# A physical model of cell metabolism

**DOI:** 10.1101/209551

**Authors:** Jorge Fernandez-de-Cossio-Diaz, Alexei Vazquez

## Abstract

Cell metabolism is characterized by three fundamental energy demands to sustain cell maintenance, to trigger aerobic fermentation and to achieve maximum metabolic rate. Here we report a physical model of cell metabolism that explains the origin of these three energy scales. Our key hypothesis is that the maintenance energy demand is rooted on the energy expended by molecular motors to fluidize the cytoplasm and counteract molecular crowding. Using this model and independent parameter estimates we make predictions for the three energy scales that are in quantitative agreement with experimental values. The model also recapitulates the dependencies of cell growth with extracellular osmolarity and temperature. This theory brings together biophysics and cell biology in a tractable model that can be applied to understand key principles of cell metabolism.

---

The basal metabolic state of a cell is characterized by a maintenance energy demand [33, 6]. It has been estimated from the extrapolation of the growth dependence of the energy demand to the zero growth limit. For mammalian cells it is particularly high, with values around 0.3 mol ATP/L/h [17]. When cells grow, move or perform other functions the energy requirements increase beyond the basal maintenance demand. Cells utilize glycolysis and oxidative phosphorylation to satisfy these energetic demands. Glycolysis has a low yield of 2 mol ATP/mol glucose [47], but it is characterized by a high horsepower (energy produced per volume of enzymes) [17, 44]. Oxidative phosphorylation has a higher yield of 32 mol ATP/mol glucose [47], but it is characterized by a lower horsepower [17, 44]. The differences in yield and horsepower imply a metabolic switch from pure oxidative phosphorylation at low energy demands to mixed oxidative phosphorylation plus obligatory fermentation (glycolysis + lactate release) at high energy demands [46, 44]. For mammalian cells this takes place at an energy demand of about 2 mol ATP/L/h [46], 10 times the energy demand of cell maintenance. Finally, there is the energy demand necessary to sustain the maximum growth rate, or a high metabolic rate in general. The maximum growth rate energy demand can only be sustained by glycolysis [44] and therefore we can estimate the maximum energy requirements of cells from their maximum reported rates of fermentation. For mammalian cells that gives us an estimate of about 8 mol ATP/L/h [46], close to an order of magnitude above the energy threshold for obligatory fermentation.

These metabolic functions are fulfilled within the context of an intracellular milieu crowded with macromolecules and organelles [43]. In fact, both the energy threshold of obligatory fermentation and maximum energy demand can be deduced from molecular crowding constraints. A good order of magnitude can be obtained using simple models focusing on energy and protein balance [4, 44]. More precise estimates can be achieved considering full scale models of cell metabolism with annotations of enzyme kinetic parameters [45, 31, 1, 32, 41, 16]. Yet, no theory has been proposed to explain the origin and magnitude of the maintenance energy demand. Here we address this problem based on the hypothesis that the maintenance energy demand corresponds to the energy expended by molecular motors to fluidize the intracellular milieu. The incorporation of this idea into a basic model of cell metabolism allows us to make quantitative predictions of the three energy scales that are in agreement with measured values. We discuss several implications of this theory to motivate further experimental work for its verification.

## Entropic pressure of molecular crowding

The machinery of life operates on the background of a gel like substance with properties distinct from ideal solutions [5]. In turn, the properties of this background substance are affected by the molecular crowding of the cellular machinery [14]. Molecular crowding hinders the diffusion of large macromolecules. When a trace particle is confined by the surrounding molecules, impeding its movement, a hole needs to be created to make space available for the trace molecule to diffuse (Fig. 1, molecule A). However, the creation of a hole lowers the entropy of the surrounding system, by reducing the number of microscopic configurations of crowding molecules and free space. Given a regular lattice of *N* spherical holes and *n* hard spheres, the entropic pressure associated with the creation of a hole of size equal to *m* spheres can be estimated as (assuming *m* ≪ *n*),

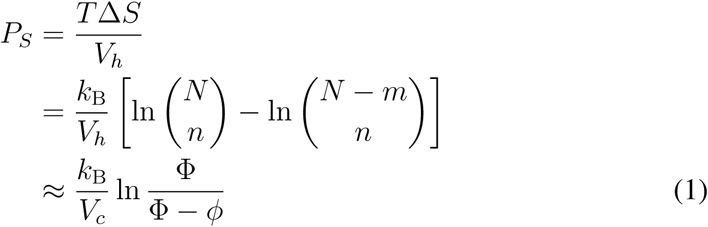

where *V*_*c*_ is the typical volume of molecular crowders (the hard spheres), *V*_*h*_ = *mV*_*c*_ the hole volume, *ϕ* = *nV*_*c*_/*V* the excluded volume fraction by the molecular crowders, Φ = *NV*_*c*_/*V* the maximum packing density and *V* the cell volume. For typical values of macromolecular volume fraction *ϕ* = 0.4 [48], maximum packing density of polydisperse spheres Φ = 0.8 [40] and protein volume *V*_*c*_ = 135nm^3^ [29], we obtain an entropic pressure of about 0.2 atm (at a temperature *T* = 25°C). This pressure is within the order of the turgor pressure in *E. Coli* [28] and, therefore, it is physiologically relevant. As expected the entropic pressure diverges as *ϕ* approaches the maximum packing density Φ, indicating that at maximum packing the creation of holes becomes unfeasible. We also note that the entropic pressure is inversely proportional to the crowder volume *V_c_*.

**Figure 1:**
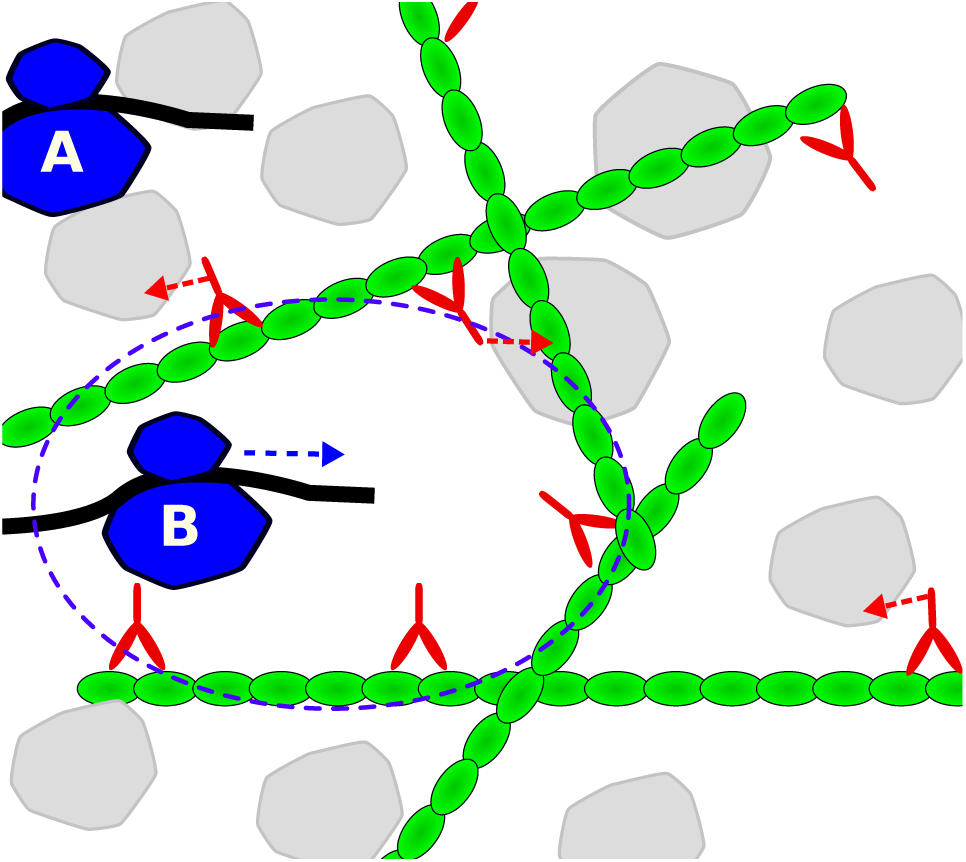
Molecular motors counteracting molecular crowding. Molecular crowders (gray) impede cellular processes such as protein synthesis. (A) A ribosome (blue) translating a mRNA strand must displace the crowders before it can move to the next codon. (B) Molecular motors (red) increase the propensity of cavities with lower density of molecular crowders where processes like translation can proceed freely.

## Molecular motors pressure

A fundamental question in cell biology is how cells can achieve high metabolic rates in the context of the highly crowded cellular environment. The experimental evidence tells us that high rates of metabolism coincide with a fluidization of the cytoplasm [34]. The fluidization of the cell milieu is most likely determined by the activity of molecular motors that push macromolecules at the expense of ATP hydrolysis [9]. This observation prompted us to the hypothesis that the maintenance energy demand may be rooted on the energy expended by molecular motors. To put this hypothesis into a working model we postulate an ideal gas of molecular motors and determine the motors pressure (Fig. 1, red molecules). As in an ideal gas, we assume the particles (motors) move in a straight line at constant speed, which is a good approximation for the movement of motors along actin filaments. The key difference is that the motors have low speeds compared to ideal gas molecules. However, motors can apply significant forces (kicks) of the order of pN. When motors get in contact with macromolecules they can transmit a large impulse (Fig. 1, arrows). Introducing this modification to the classical kinetic theory of gases we obtain the motors pressure (Supplementary Text),

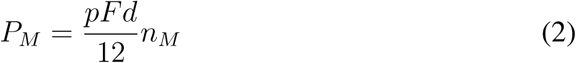

where *F* and *d* are the motors force and displacement per kick, *n*_*M*_ is the number of molecular motors per cell volume, and *p* is the motors persistence, a non-dimensional parameter quantifying the motors tendency to maintain their direction of motion upon contact with macromolecules.

The cell could tune the motors concentration in order to counteract the entropic pressure of molecular crowding. The optimal motors concentration should be a balance between the tendency of molecular motors to open holes and facilitate movement (Fig. 1, molecule B) and the energetic cost of motors activity. We postulate that the optimal solution occurs when the motors pressure matches the entropic pressure of molecular crowding *P*_*M*_ = *P*_*S*_. From this postulate and equations (1) and (2) we obtain the volume fraction occupied by molecular motors, *ϕ*_*M*_ = *V*_*M*_^*n*^_*M*_,

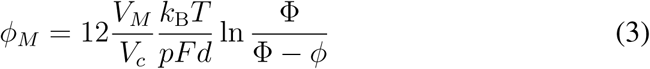

where *V*_*M*_ is the typical volume of molecular motors. Within this framework, the volume fraction of molecular motors is proportional to the ratio of the thermal energy *k*_B_*T* to the motors energy per kick *Fd*. It is also evident that the volume fraction of molecular motors is a monotonically increasing function of the macromolecular fraction *ϕ*.

## Basic model of cell metabolism

Now we are ready to investigate a model of cell metabolism where the maintenance energy demand represents the energy expended by molecular motors to counteract molecular crowding. Since proteins are the major component of cell biomass, we will focus on protein metabolism for the sake of simplicity. We divide proteins/organelles in five classes based on their function. Background proteins (with volume fraction *ϕ*_0_), ribosomes (*ϕ*_*R*_), molecular motors (*ϕ*_*M*_), fermentation enzymes (*ϕ*_*F*_) and oxidative phosphorylation machinery (*ϕ*_0_). We further assume that cells are in an exponential growth phase, with growth rate *µ*, and that they have an excess supply of carbon sources for energy production and amino acids for protein synthesis. This simplified model is governed by the equations,

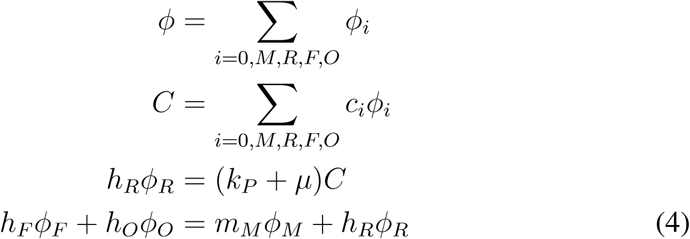

where *i* = 0*, M, R, F, O* is the compartment index, *C* the protein concentration in the cell (mol of amino acids/cell volume), *c*_*i*_ the protein concentration in compartment *i* (mol of amino acids/compartment volume), *k*_*P*_ the protein turnover rate (amino acids/unit of time), *h*_*R*_ the ribosome horsepower (moles of amino acid added to new proteins/volume of ribosome/unit of time), *h*_*F*_ and *h*_*O*_ the fermentation and oxidative phosphorylation horsepowers (moles of ATP/volume of enzyme/unit of time), *m*_*M*_ the maintenance energy rate per motor volume (moles of ATP consumed/volume of motor/unit of time), and *e*_*P*_ the energy demand of protein synthesis (molecules of ATP consumed per amino acid). The first two equations in (4) are composition equations, encoding the proper bookkeeping of volume and protein. The last two equations are metabolic balance equations for proteins and energy production/consumption.

The system of equations (4) can be solved to determine the growth rate. The solution can be written in a compact form (see Supplementary text),

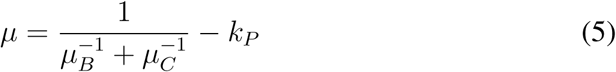

where

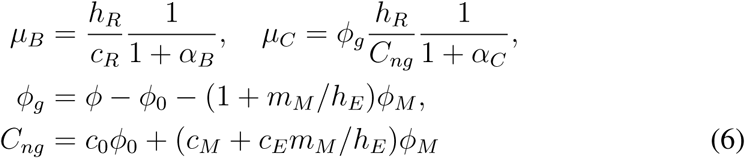

and *α*_*B*_ = *e*_*P*_(*c*_*E*_*h*_*R*_)/(*c*_*R*_*h*_*E*_) and *α*_*C*_ = *e*_*P*_ *h*_*R*_/*h*_*E*_ are the biosynthetic and molecular crowding costs, respectively. Equation (5) indicates that the growth rate is the harmonic mean between the biosynthetic and molecular crowding growth rates. The biosynthetic growth rate (*µ*_*B*_), when neglecting molecular crowding, is basically dictated by the ribosome efficiency *h*_*R*_/*c*_*R*_(amino acids added to new proteins per ribosome). The molecular crowding growth rate (*µ*_*C*_), when neglecting the biosynthesis cost, is proportional to the volume fraction of growth-associated components *ϕ*_*g*_ and the ribosomes horsepower *h*_*R*_ and it is inversely proportional to the growth-independent component of the protein content *C*_*ng*_. The biosynthetic and crowding costs *α*_*B*_ and *α*_*C*_ are corrections taking into account the protein content and volume fraction, respectively, that are not available to the ribosomes because of they are reserved for the energy producing machinery. The index *E* stands for the energy generation pathway, including pure fermentation *E* = *F*, pure oxidative phosphorylation *E* = *O*, or mixed fermentation/oxidative phosphorylation.

## Osmolarity dependency of the growth rate

Figure 2A shows the dependency of the growth rate with the excluded volume fraction *ϕ*, for the case when energy is produced by pure oxidative phosphorylation (blue line) or fermentation (black line). At any given *ϕ* the fermentation line exhibits a higher value, indicating that fermentation can sustain higher growth rates than oxidative phosphorylation. This is explained by the fact that fermentation has a higher horsepower (*h*_*F*_ > *h*_*O*_), *i.e.* it can produce more energy per volume of enzyme than oxidative phosphorylation. We also note that the plot of *µ* vs. *ϕ* follows a bell like shape. A similar behavior has been observed for the growth rate dependency with medium osmolarity [10, 39, 7, 23, 36, 27]. The ideal osmotic response of cells from a reference state (^∗^) to a new external osmolarity follows the Boyle–van’t Hoff relation Π(*V*− *b*) = Π^∗^(*V* ^∗^ − *b*) [25], where Π^∗^ and Π are the osmotic pressures at the reference and new states, *V* ^∗^ and *V* are the corresponding cell volumes, and *b* is the solvent excluded volume occupied by macromolecules. *b* is assumed to be constant between the two conditions as observed experimentally. The Boyle–van’t Hoff relation can be rewritten to obtain a relationship between *ϕ* = *b/V* and the osmotic pressure, *ϕ* = Π*/*(Π_50_ + Π), where Π_50_ = Π^∗^*ϕ*^∗^*/*(1 –*ϕ*^∗^) is the osmotic pressure where macromolecules occupy 50% of the cell volume. Substituting latter equation into the growth rate equation (5) we can plot the growth rate as a function of the relative osmotic pressure Π*/*Π_50_ (Figure 2B). The resulting behavior explains the observed bell shape plot of growth rate vs. osmolarity.

**Figure 2:**
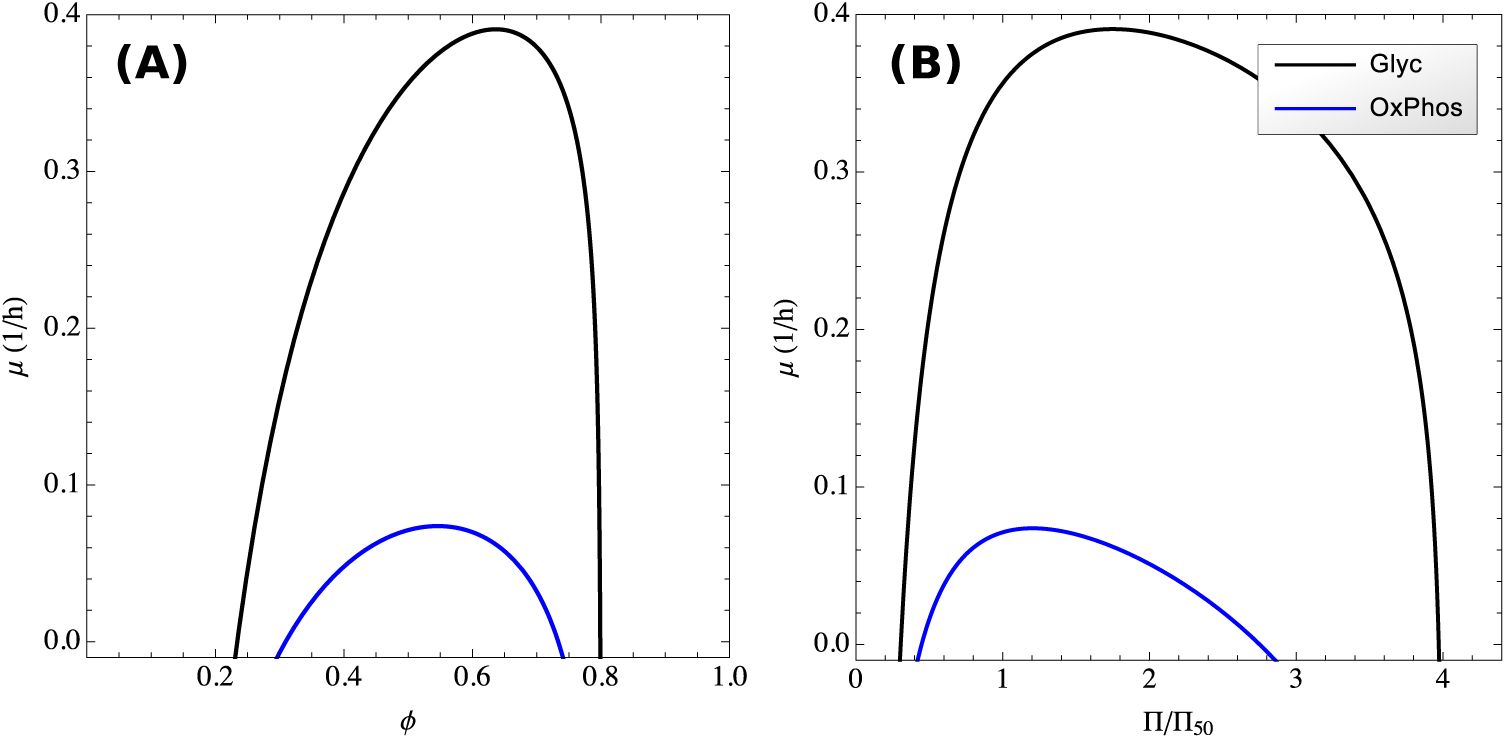
Osmolarity dependency of the growth rate. (A) Maximum growth rate as a function of the excluded volume fraction, when energy is produced by fermentation (black line) or oxidative phosphorylation (blue line). (B) Maximum growth rate as a function of external osmolarity. These plots were obtained using parameters for mammalian cells.

## Energy scales of cell metabolism

At a given growth rate, the total energy demand and maintenance energy demand can vary significantly depending of the excluded volume fraction (Fig. 3A,B). However, solutions that are further constrained to satisfy some optimality criteria (*e.g.*, maximum yield or minimum carbon consumption) are concentrated on the lower *ϕ* range of the feasible space. As expected the overall energy demand increases with increasing the growth rate for the optimal solutions (Fig. 3A). In contrast, it is not clear *a priori* whether the maintenance energy demand changes or not with the growth rate. From equation (3) it follows that solutions where *ϕ* is constrained to be a constant are characterized by a constant *ϕ*_*M*_ and, therefore, a growth independent maintenance energy demand. We also take a closer look to the maintenance energy demand for the solution maximizing growth yield (Fig. 3B, orange line). In this case, the maintenance energy demand remains approximately constant relative to the total energy demand, except for the extreme values close to zero and maximum growth rate. This implies that the energy of cell maintenance may appear to be independent of the growth rate even though we have not imposed such a constraint.

**Figure 3:**
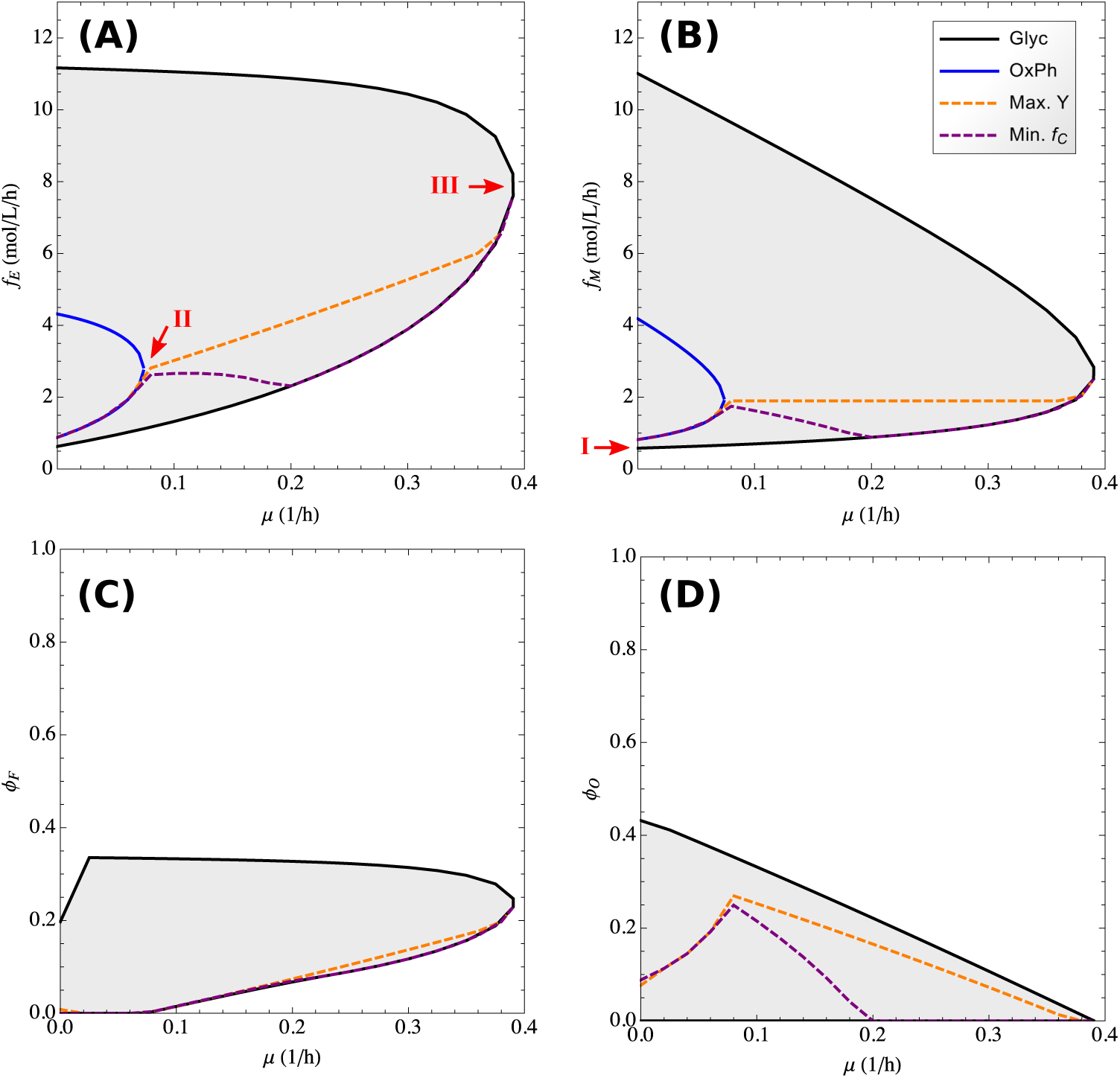
Energy balance as a function of growth rate. (A,B) Energy demand of all metabolism (A) and cell maintenance (B) as a function of the growth rate. The arrows indicate the three energy scales of metabolism: I. maintenance, II. switch to fermentation, and III. maximum growth rate. (C,D) Volume fractions of fermentation enzymes (C) and oxidative phosphorylation machinery (D) as a function of growth rate. The shaded area represents the range of feasible values corresponding to different values of *ϕ*. The dashed lines show the trajectories corresponding to lowest carbon consumption (purple) and maximum growth yield (orange). These plots were obtained using parameters for mammalian cells.

An important question in cell metabolism is why cells exhibit active fermentation even under conditions where oxygen is present and oxidative phosphorylation should be the pathway of choice. This problem can be addressed within our model by analyzing its solution when both fermentation and oxidative phosphorylation can be active. Because of their redundant nature for energy generation, the volume fractions occupied by the oxidative phosphorylation and fermentation machineries vary widely (Fig. 3C and D). Since the maximum growth rate sustained by oxidative phosphorylation is smaller than that sustained by fermentation, it is evident that there is a threshold growth rate above which fermentation is obligatory (Fig. 3C). In compensation the maximum rate of oxidative phosphorylation goes down with increasing proliferation rate (Fig. 3D). The later behavior is in agreement with the observed decrease of mitochondria protein mass in yeast [27] and mammalian cells [46] at higher growth rates.

Taking all this evidence together we provide an explanation for the three key energy scales of cell metabolism. From the quantitative point of view we focus on values for mammalian cells. The maintenance energy demand is represented by the energy expend by molecular motors. Provided that the motors persistence parameter *p* is of the order of 1, the energy demand of cell maintenance predicted from this model (~ 0.5 mol/L/h, Fig. 3B, I) is in the range of what is observed experimentally (around 0.3 mol ATP/L/h [17]). This is a striking observation given that our estimate is based on microscopic parameters characterizing molecular motors (Methods). The energy demarking the switch to obligatory fermentation is simply a consequence of fermentation having a larger horsepower than oxidative phosphorylation. This result in an energy demand to switch to obligatory fermentation of ~3 mol/L/h (Fig. 3A, II) in the range of experimental observations (~ 2 mol ATP/L/h [46]). Finally, the energy demand at maximum growth rate is simply determined by the energy demand at the maximum growth rate that can be achieved using fermentation (~ 8 mol/L/h, Fig. 3A, III), again in very good agreement with the maximum fermentation rates reported for mammalian cells (~ 8 mol ATP/L/h [46]).

## Background proteins reduce the metabolic capacity

Another important observation is the dependency of the model solution with the volume fraction of background proteins *ϕ*_0_. The solutions described above were obtained assuming a typical value of *ϕ*_0_ = 0.2 as observed for mammalian cells (Methods). However, changing *ϕ*_0_ can have a big quantitative impact on the model solution. Reducing *ϕ*_0_ by half we obtain a significant increase in the maximum growth rate and the range of *ϕ* with feasible solutions (Fig. 4). In contrast, doubling *ϕ*_0_ reduces the maximum growth rate and constraints the solutions to a smaller *ϕ* range concentrated near the maximum packing density (Fig. 4). For a healthy organism, we think there is not much flexibility on the *ϕ*_0_ values. First, the background protein content is constrained by the actin cytoskeleton that is required to support the molecular motors activity. Just to get an order of magnitude, the rate of ATP hydrolysis by myosin is characterized by a half saturation constant for actin of the order of 40 *µ*M [42, 2]. That value multiplied by a typical molar mass of 500 kDa and a specific excluded volume of 2 mL/g yields a volume fraction of 0.04. On top of that we need to add the volume of the nuclei (eukaryotes) or nucleoid regions (prokaryotes). The nucleus occupies a volume fraction of 0.06 to 0.1 in mammalian cells [3, 29, 28], bringing the lower bound of *ϕ*_0_ to about 0.15. Third, there are other metabolic enzymes associated with lipid and nucleotide metabolism that can occupy a significant volume fraction. For example, fatty acid synthase alone occupies a typical volume fraction of 0.007 (Methods). Therefore, a value of *ϕ*_0_ = 0.2 is about what we expect for mammalian cells. However, *ϕ*_0_ takes different values in other organisms. For yeast cells *ϕ*_0_ ≈ 0.1, and that may explain why yeast cells can grow faster than mammalian cells (Fig. 4, *ϕ*_0_ = 0.1 vs 0.2). It is also worth noticing that *ϕ*_0_ could increase above 0.2 in pathological conditions where protein aggregation occurs. In such instances there can be a dramatic reduction in the metabolic capabilities of cells (Fig. 4, *ϕ*_0_ = 0.4 vs 0.2).

**Figure 4:**
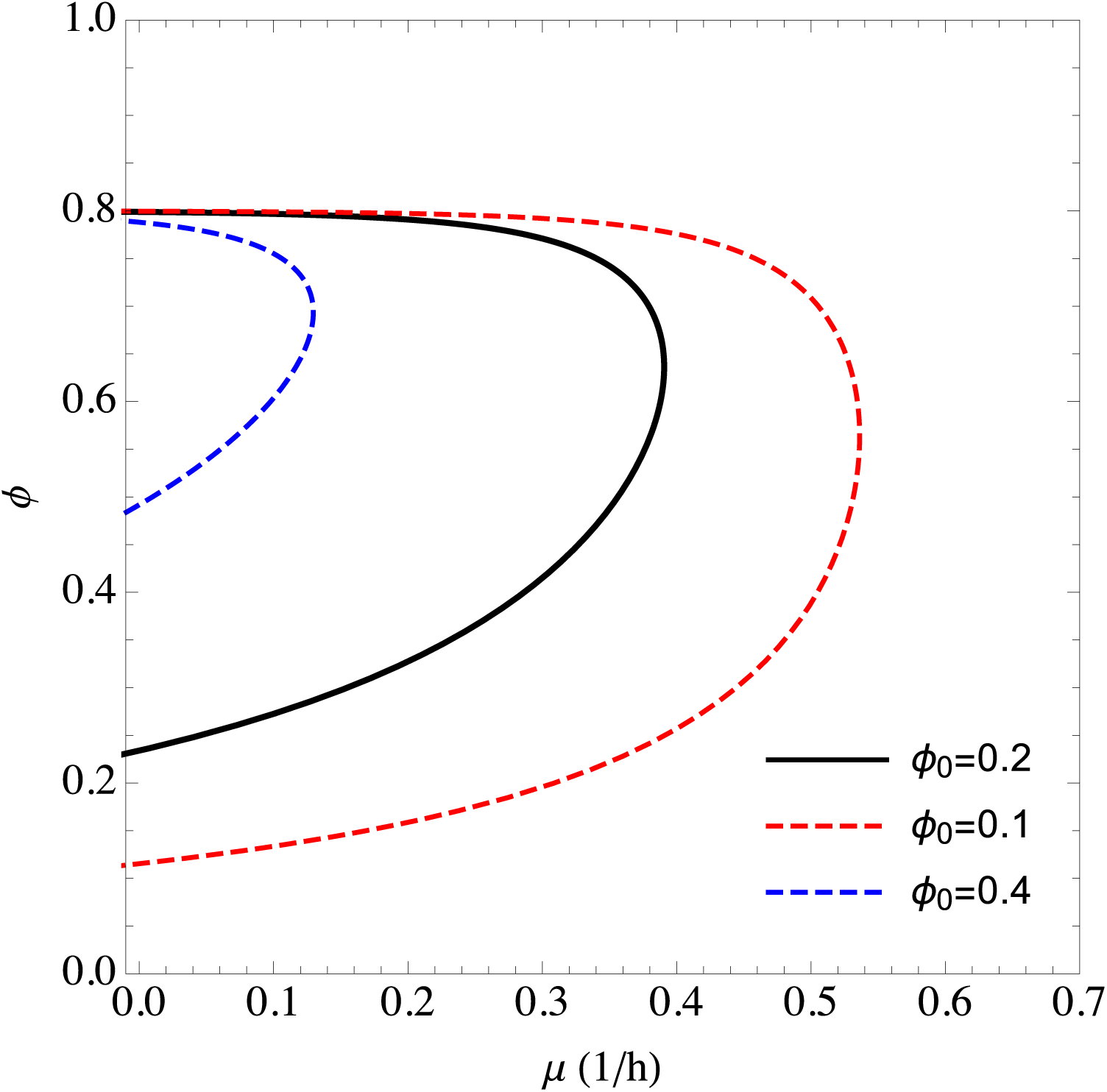
Impact of the volume fraction of background proteins. The space of feasible solutions in the *µ, ϕ* plane for different values of *ϕ*_0_.

## Temperature dependency of the growth rate

The molecular motors fraction that is required to counteract molecular crowding is proportional to the temperature (3). This implies that the cost associated with molecular motors is higher with increasing temperature and should result in a drop of metabolic rate at high temperatures. That together with the Arrhenius-like increase of enzyme rates with temperature, *e*^−*E*_*a*_/*k*_*B*_*T*^, would result in a maximum growth rate at some intermediate temperature. Figure 5 shows the temperature dependence of the growth rate, after accounting for the Arrhenius temperature dependencies of the horsepower of ribosomes and fermentation and the kicking rate of molecular motors. The qualitative shape of the curve is strikingly similar to what is observed experimentally [30]. The precise temperature where the maximum growth rate is achieved is in part determined by the actual values of the activation energies. It is quite impressive that this simple model can reproduce the temperature dependency of the growth rate without taking into consideration the propensity of proteins to denature at high temperatures.

**Figure 5:**
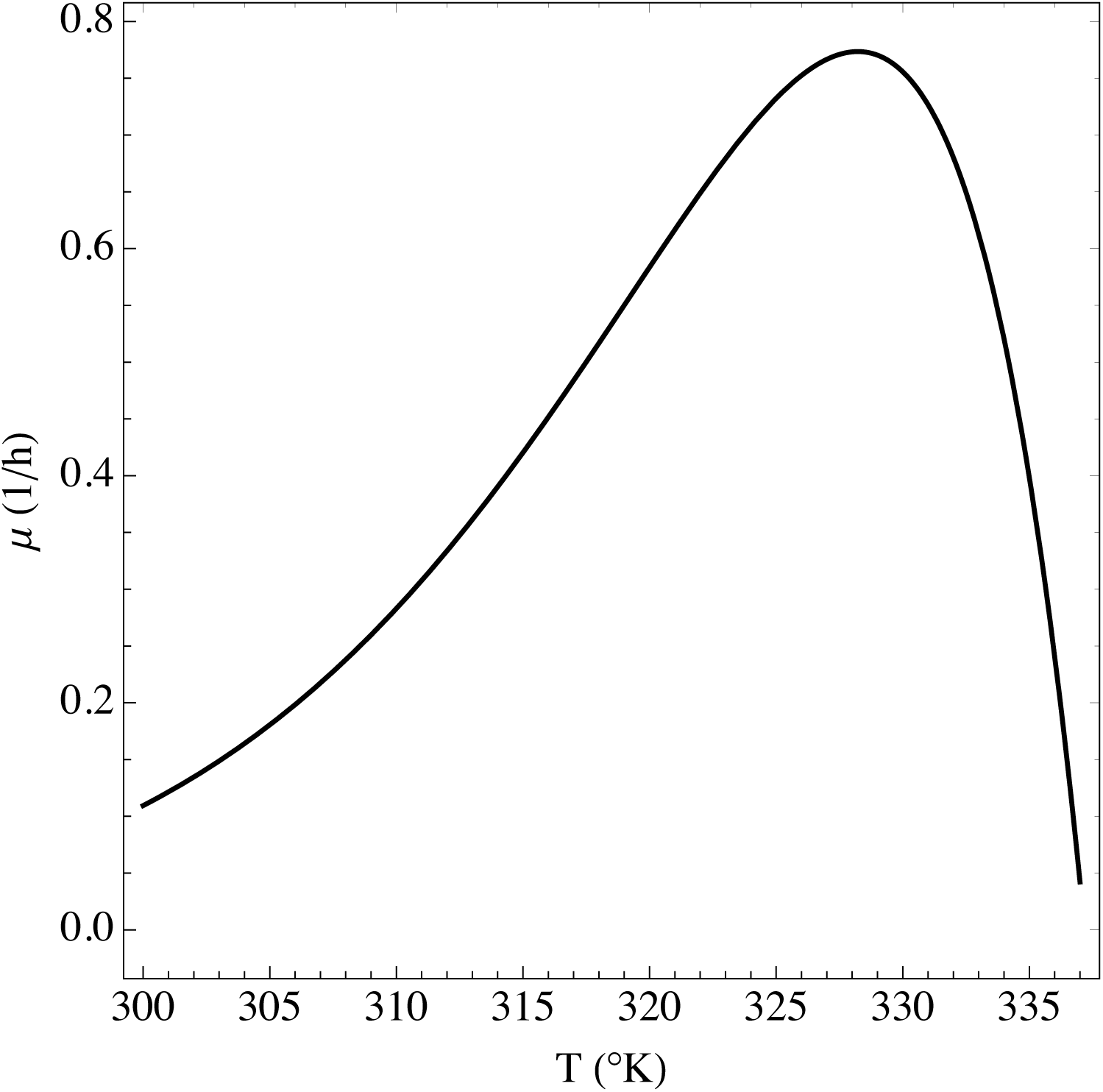
Temperature dependency of the maximum growth rate. This plot was obtained using parameters for mammalian cells.

## Conclusions

We have investigated a physical model of cell metabolism where the energy demand of cell maintenance represents the energy expended by molecular motors to counteract the entropic forces associated with molecular crowding. The model predicts an energy demand for cell maintenance that is in the range of what is observed experimentally. More importantly, the model exhibits a rich behavior depending on the macromolecular volume fraction and growth rate. It predicts the growth rate threshold for the metabolic switch to obligatory fermentation and the maximum growth rate. The model also explains the bell shape curve of growth rate as a function of osmolarity and temperature.

There are several predictions that will require further testing for verification. The model predicts that the maintenance energy demand is not necessarily independent of the growth rate. Therefore we cannot assume a linear extrapolation of the energy demand to the zero growth rate. Based on our analysis this would be valid only if cells operate at constant macromolecular volume fraction. There are however other scenarios where the maintenance energy demand is a function of the growth rate, even through it may appear linear for a certain range of growth rates.

Further work is also required to understand the impact of protein denaturation. The increasing demand of molecular motors at higher temperatures together with the Arrhenius dependency of reaction rates is sufficient to explain the existence of an optimal temperature where growth rate is maximum. Protein denaturation will introduce further corrections due to a reduction of enzymes efficiency and due to the formation of protein aggregates. Which of these factors dominate is not clear at this point.

It is also evident that a metabolic switch to obligatory fermentation can be obtained in many ways. We have discussed the metabolic switch with increasing the growth rate. The same behavior can be achieved by increasing the rate of protein turnover, effectively modeling the impact of protein secretion on cell metabolism. That may explain why non-growing fibroblasts with high rates of protein secretion exhibit high rates of fermentation [24]. A metabolic switch to obligatory fermentation could be also achieved at constant growth rate and going from low to high macromolecular excluded volume fraction.

Finally, the model introduced here can be extended to more comprehensive representations of cell metabolism. To this end we can introduce the more general formulation of metabolic balance *Sf* = 0, with *f* representing the vector of steady state reaction rates and *S* the stoichiometric matrix of the metabolic network. These metabolic fluxes would be further constrained by the reaction rate capacities *f*_*i*_ ≤ *h*_*i*_*ϕ*_*i*_, with *h*_*i*_ representing reaction horsepowers and *ϕ*_*i*_ enzyme volume fractions. This generalization will allow us to understand the impact of molecular crowding and the counteracting motors activity on metabolic pathways beyond energy and protein metabolism.

## Methods

### Parameter estimation

#### Proteins specific excluded volume

The average specific excluded volume of macromolecules in cells is *v*_*s*_ = 2 mL/g for proteins (Figure A2c in Ref. [48]). Since cells are made mostly made of proteins, 2 mL/g can be taken as an estimate of the typical specific excluded volume of proteins in cells. An independent estimate has been obtained by electrospray ionization mass spectrometry, which results in an average effective protein density of 0.58 g/mL [21], or equivalently a specific excluded volume of 1.7 mL/g, close to the 2 mL/g value reported for cell extracts.

#### Volume fraction of fatty acid synthase

Using publicly available data [19], we calculated an average mass fraction of fatty acid synthase in the proteome of the NCI60 panel of cancer cell lines of 0.02. Multiplying by the average protein density of 0.15 g/mL [13] and dividing by the protein excluded volume, 2 mL/g, yields the estimated volume fraction of 0.007.

#### Volume fraction of background proteins

Following the same procedure as in the estimation of the volume fraction of fatty acid synthase, we estimated the volumetric fraction of background proteins by computing the fraction left after excluding fermentation enzymes, molecular motors, mitochondrial and ribosome proteins. Protein localization and function were distinguished following annotations from the Gene Ontology database [12]. The estimated volume fraction occupied by background proteins varies from 0.1 up to 0.4 in the panel of NCI60 cell lines [19, 13], with an average value of 0.2.

#### Compartmental protein concentrations

The compartmental protein concentrations of fermentation, motors and background proteins was calculated using *c*_*i*_ = *m*_*a*_/*v*_*s*_ = 4.6 mol/L, where *i*= *F, M,* 0, *m*_*a*_ = 109 g/mol is the average molar mass of amino acids and *v*_*s*_ = 2 mL/g is the protein specific excluded volume. Using the eukaryotic ribosome volume of 4000 nm^3^ and a composition of 11590 amino acids/ribosome [26, 22], we calculated *c*_*R*_ = 4.8 mol/L. The specific volume of mitochondria in mammalian cells is ~ 2.6 mL of mitochondria/g of mitochondrial protein. Multiplying by *m*_*a*_ we obtain *a*_*O*_ ≈ 3.5 mol/L.

#### Fermentation horsepower

Reference [38] reports the rate of fermentation between 0 and 40C from an *in vitro* reconstitution of glycolysis enzymes with 40 mg/mL of total glycolytic protein concentration. This data is well fitted to the Arrhenius law of the rate of a biochemical reaction *f*_*F*_ = *f*_0_*e*^−*E*_*a*_/*RT*^, with *f*_0_ = 33 *µ*mol/min/mL and *E*_*a*_ = − 77 kJ/mol. Using this Arrhenius law we obtain the fermentation rate at 37°C to be 47 *µ*mol/min/mL. Dividing by a protein excluded volume of 2 mL/g we obtain a fermentation horsepower of *h*_*F*_ = 35 mol ATP/L/h. A theoretical estimate can be obtained under the assumption that all enzymatic steps of glycolysis are at saturation, resulting in *h*_*F*_ = 1/*v*_*s*_ Σ_*i*_(*s*_*i*_/*k*_*i*_), where *v*_*s*_ is the proteins specific excluded volume, the sum run over all glycolysis enzymes, *s*_*i*_ is the stoichiometric coefficient of reaction *i* relative to lactate release and *k*_*i*_ is the specific turnover rate of the enzyme catalyzing step *i*. Based on specific turnover data from BRENDA [37], *k* = 36.5, 620, 90, 22.3, 8000, 68.1, 600, 1.5, 70, 400, and 106 *µ*mol/mg/min for the enzymes hexokinase, phosphoglucose isomerase, phosphofructose kinase, alodolase, glyceraldehyde-3-phosphate dehydrogenase, phosphoglycerate kinase, phosphoglycerate mutase, enolase, pyruvate kinase and lactate dehydrogenase, respectively. Furthermore, *s*_*i*_ = 1/2 from the hexokinase to the aldolase step and *s*_*i*_ = 1 from glyceraldehyde-3phosphate dehydrogenase to lactate dehydrogenase. Using these enzyme specific turnovers, stoichiometric coefficients, and a protein specific excluded volume of 2 mL/g we obtain *h*_*F*_ = 40 mol ATP/L/h. This theoretical estimate is in excellent agreement with the experimentally measured value. The calculations and figures reported in the main text were obtained using the experimental value of *h*_*F*_ = 35 mol ATP/L/h.

#### Oxidative phosphorylation horsepower

The oxidative phosphorylation horsepower in mammalian cells can be estimated from experimental reports of the maximum capacity for ATP production by isolated mitochondria [44]. The average value is *h*_*O*_ = 10 ATP mol/L of mitochondria/h for mitochondria isolated from healthy mammalian cells. For mitochondria isolated from cancer cells it goes down to 3 mol ATP/L of mitochondria/h. A theoretical estimate was obtained using a mathematical model of mitochondrial oxidative phosphorylation [20]. Constraining the total enzymatic mass of oxidative phosphorylation and then maximizing the rate of ATP production, we obtain a theoretical oxidative phosphorylation horsepower of 19 mol ATP/L of mitochondria/h. This value closely matches the maximum mitochondrial horsepower reported across multiple cell types [44]. The calculations and figures reported in the main text were obtained using the average experimental report for healthy mitochondria of *h*_O_ = 10 ATP mol/L of mitochondria/h.

#### Ribosomes horsepower

The horsepower of mammalian ribosomes is *h*_*R*_ = 8.4 mol of amino acid incorporated/L of ribosome/h [44].

#### Motors parameters

Molecular motors exert forces of the order of *F* = 5 pN per kick moving for about *d* = 10 nM [11, 18]. The motors kicking rate is ~*κ* = 5*s*^−1^, while the duration of a kick is *τ* ~ 0.5 s [35, 29]. This results in a persistence parameter, *p* = *rκτ*, where *r* is the number of repeated kicks upon contact with a macromolecule. The average repeated kicks *r* ≈ 5 was estimated so as to match a maintenance demand of ≈ 1 mol/L/h at a typical macromolecular volume fraction of *ϕ* = 0.4 [48]. The motors maintenance energy per motor volume was calculated as *m*_*M*_ = *κ/*(*M*_*M*_ *v*_*s*_), where *M*_*M*_ is the motors molar mass and *v*_*s*_ is the protein specific excluded volume. We use the molar mass of Myosin Va, *M*_*M*_ = 215405 g/mol (UNIPROT:Q9Y4I1) as a typical value. This results in *m*_*M*_ = 42 mol/h/L.

#### Activation energies

The biochemical horsepowers were assumed to follow an Arrhenius dependence on the temperature, 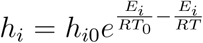, where *i* = *R,F,T*_0_ is the reference temperature where the horsepower equals *h*_*i*0_ and *E*_*i*_ is the activation energy. Since the motors kicking is also an activated process, the maintenance requirement of motors was assumed to follow a similar law, 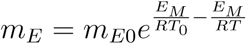. The activation energies were estimated from the literature: ribosomes 80 kJ/mol [15], glycolysis 77 kJ/mol [38], and myosin 125 kJ/mol [8].

## Acknowledgements

This work was supported by Cancer Research UK C596/A21140. This project has received funding from the European Unions Horizon 2020 research and innovation programme MSCA-RISE-2016 under grant agreement No. 734439 INFERNET.

## Author contributions

JFC and AV developed the theory and wrote the manuscript.

